# Phylogenetic relationships among haloalkaliphilic archaea of the family *Natrialbaceae*

**DOI:** 10.1101/2020.01.20.913392

**Authors:** Shivakumara Siddaramappa

**Affiliations:** Institute of Bioinformatics and Applied Biotechnology, Biotech Park, Electronic City, Bengaluru - 560100, Karnataka, INDIA

**Keywords:** Halobacteria, Natrialbales, Natrialbaceae, Natrialba, Proteome, Phylogeny

## Abstract

The family *Natrialbaceae* is a member of the class *Halobacteria* of the archaeal phylum *Euryarchaeota*. Seventeen genera with validly or effectively published names are currently included within this family. In this study, using pairwise average nucleotide identity and average amino acid identity comparisons in conjunction with phylogenetic analysis, it has been shown that the family *Natrialbaceae* is highly diverse and contains several potentially novel species and genera that are yet to be fully characterized. The deduced proteome sequence-based phylogenetic tree, constructed using the alignment- and parameter-free method CVTree3, contained six major clades, with *Salinarchaeum* sp. Harcht-Bsk1 being the only representative within clade 1. Furthermore, *Haloterrigena daqingensis* was found to be closely related to *Natronorubrum sediminis*, and it is proposed that these archaea together represent a novel genus. Interestingly, *Haloterrigena jeotgali*, *Haloterrigena thermotolerans*, and *Natrinema pellirubrum* were found to be very closely related to each other, and it is proposed that they be merged into a single species. Notably, the type genus *Natrialba* itself appeared to be heterogenous and contains species that could be broadly classified among two genera. Likewise, the genus *Natrinema* is also heterogenous and contains species that could be classified among six genera. Altogether, 19 novel genera have been proposed to be created, and four haloalkaliphilic archaea hitherto recognized only using genus names are confirmed to represent novel species.

## INTRODUCTION

Woese *et al*. (1990) subdivided the domain Archaea into two kingdoms and named them as *Euryarchaeota* and *Crenarchaeota*. The formal name *Euryarchaeota* reflected the variety of metabolic features of, and the different types of niches occupied by, the methanogenic archaea that were included in this kingdom. Within the hierarchical system of the second edition of the Bergey’s manual of systematic bacteriology, these two kingdoms became two novel phyla of archaea (Garrity and Holt 2001a, 2001b). When it was proposed, the phylum *Euryarchaeota* contained seven novel classes, including *Halobacteria* (Garrity and Holt, 2001b). This class contained a single order (*Halobacteriales*), which had a single family (*Halobacteriaceae*). The novel genus *Natrialba* was proposed by Kamekura and Dyall-Smith (1995), and was among the several genera that were included in the family *Halobacteriaceae* within the the Bergey’s manual (Garrity and Holt, 2001b). Using phylogenetic analyses, Gupta *et al*. (2015) proposed a novel order (*Natrialbales*) containing a novel family (*Natrialbaceae*) within the class *Halobacteria*. When it was circumscribed, the family *Natrialbaceae* contained 12 genera, of which six had been proposed since 2002 (Gupta *et al*., 2015). The analyses by Gupta *et al*. (2015) relied on 16S rDNA gene sequences and concatenated sequences of 32 conserved proteins. Recently, Chun *et al*. (2018) proposed a set of minimal standards for prokaryotic taxonomy, and opined that “a multigene-based phylogenomic treeing approach should be the choice for defining genera or higher taxa”. With the availability of the genome sequences of several new genera/species of haloarchaea in the last five years, a comprehensive ‘phylogenomic treeing approach’ seemed feasible. In this context, the objectives of the present study were to perform pairwise comparisons of whole genome/proteome sequences of various genera/species of *Natrialbaceae*, and use the pairwise comparisons in conjunction with phylogenetic analysis to decipher the taxonomic relationships among archaea of this family.

## MATERIALS AND METHODS

### Genome and deduced proteome sequence comparisons

At the time of writing (July 2019), the National Center for Biotechnology Information (NCBI) taxonomy browser (https://www.ncbi.nlm.nih.gov/Taxonomy/Browser/wwwtax.cgi?mode=Undef&id=1644061&lvl=3&lin=f&keep=1&srchmode=1&unlock) had listed 17 genera within the family *Natrialbaceae*. Genome sequences of these archaea were obtained from GenBank using the links within the taxonomy browser. Deduced proteome sequences were obtained from the UniProt database (https://www.uniprot.org/proteomes/?query=Natrialbales&sort=score). The final list included 67 genome (and their deduced proteome) sequences from 60 strains (Table 1). Forty eight of these were type strains of different species (Table 1), of which 12 were type species of various genera. The 60 strains were from the genera *Halobiforma* (Hezayen *et al*., 2002), *Halopiger* (Gutiérrez *et al*., 2007), *Halostagnicola* (Castillo *et al*., 2006b), *Haloterrigena* (Ventosa *et al*., 1999), *Halovivax* (Castillo *et al*., 2006a), *Natrarchaeobius* (Sorokin *et al*., 2019a), *Natrialba* (Kamekura & Dyall-Smith, 1995), *Natrinema* (McGenity *et al*., 1998), *Natronobacterium* (Tindall *et al*., 1984), *Natronococcus* (Tindall *et al*., 1984), *Natronolimnobius* (Itoh *et al*., 2005), *Natronorubrum* (Xu *et al*., 1999), and *Salinarchaeum* (Dominova *et al*., 2013). Since genome and/or deduced proteome sequences were not available for any of the strains assigned to the genera *Halovarius* (Mehrshad *et al*., 2015), *Natribaculum* (Liu *et al*., 2015), *Natronobiforma* (Sorokin *et al*., 2018), and *Saliphagus* (Yin *et al*., 2017), they could not be included in the analysis. Of the 67 genome sequences, 16 were complete and 51 were draft. Two-way average nucleotide identity (ANI) and average amino acid identity (AAI) were calculated using the Kostas lab web server (http://enveomics.ce.gatech.edu) with default parameters.

**Table 1.** List of archaea whose genome sequences were compared, and deduced proteome sequences were used for phylogenetic analysis

| Serial No. | Species | Strain | Taxonomy ID | Genome accession | Proteome ID | Predicted proteins | Reference | Subclade in Fig. 1 | Emendation/ Proposal |
| --- | --- | --- | --- | --- | --- | --- | --- | --- | --- |
| 1 | *Salinarchaeum* sp. | Harcht-Bsk1 | 1333523 | CP005962 | UP000014072 | 3013 | Dominova *et al.* (2013) | 1 | None |
| 2 | *Halovivax asiaticus* | JCM 14624<sup>T</sup> | 1227490 | AOIQ00000000 | UP000011560 | 3131 | Castillo *et al.* (2006a) | 2 | None |
| 3 | *Halovivax ruber* | DSM 18193<sup>T</sup> | 797302 | CP003050 | UP000010846 | 3099 | Castillo *et al.* (2007) | 2 | None |
| 4 | *Halostagnicola larsenii* | XH-48<sup>T</sup> | 797299 | CP007055 | UP000019024 | 2843 | Castillo *et al.* (2006b) | 3 | None |
| 5 | *Halostagnicola kamekurae* | DSM 22427<sup>T</sup> | 619731 | FOZS00000000 | UP000199199 | 4009 | Nagaoka *et al.* (2010) | 3 | None |
| 6 | *Halostagnicola* sp. | A56 | 1495067 | JMIP00000000 | UP000027027 | 2711 | Kanekar *et al.* (2015) | 3 | sp. nov. |
| 7 | *Natronococcus occultus* | SP4<sup>T</sup> | 694430 | CP003929 | UP000010878 | 3888 | Tindall *et al.* (1984) | 4 | None |
| 8 | *Natronococcus amylolyticus* | DSM 10524<sup>T</sup> | 1227497 | AOIB00000000 | UP000011688 | 4317 | Kanal *et al.* (1995) | 4 | None |
| 9 | *Natronococcus jeotgali* | DSM 18795<sup>T</sup> | 1227498 | AOIA00000000 | UP000011531 | 4417 | Roh *et al.* (2007) | 4 | None |
| 10 | *Natrialba asiatica* | ATCC 700177<sup>T</sup> | 29540 | AOIO00000000 | UP000011554 | 4175 | Kamekura and Dyall-Smith (1995) | 5A | None |
| 11 | *Natrialba taiwanensis* | DSM 12281<sup>T</sup> | 1230458 | AOIL00000000 | UP000011648 | 4374 | Hezayen *et al.* (2001) | 5A | None |
| 12 | *Natrialba aegyptia* | DSM 13077<sup>T</sup> | 1227491 | AOIP00000000 | UP000011591 | 4399 | Hezayen *et al.* (2001) | 5A | None |
| 13 | *Natrialba chahannaoensis* | JCM 10990<sup>T</sup> | 1227492 | AOIN00000 000 | UP000011693 | 4030 | Xu *et al.* (2001) | 5A | gen. nov. along with serial numbers 15, 16, and 17 |
| 14 | *Natrialba magadii* | ATCC 43099<sup>T</sup> | 547559 | AOHS01000 000 | UP000011543 | 4196 | Kamekura *et al.* (1997) | 5A |  |
| 15 | *Natrialba magadii* | ATCC 43099<sup>T</sup> | 547559 | CP001932 | UP0000001879 | 3556 | Kamekura *et al.* (1997) | 5A |  |
| 16 | *Natrialba hulunbeirensis* | JCM 10989<sup>T</sup> | 1227493 | AOIM00000 000 | UP000011519 | 3852 | Xu *et al.* (2001) | 5A |  |
| 17 | *Natrialba* sp. | SSL1 | 1869245 | MASN0000 0000 | UP000179897 | 4124 | None | 5A | gen. nov. sp. nov. |
| 18 | *Natronolimnobius baerhuensis* | CGMCC 1.3597<sup>T</sup> | 253108 | MWPH0100 0000 | UP000196084 | 3662 | Itoh *et al.* (2005) | 5B | None |
| 19 | *Halopiger xanaduensis* | DSM 18323<sup>T</sup> | 797210 | CP002839 | UP0000006794 | 3588 | Gutiérrez *et al.* (2007) | 5B | None |
| 20 | *Halopiger aswanensis* | DSM 13151<sup>T</sup> | 148449 | RAPO00000 000 | UP000283805 | 4631 | Hezayen *et al.* (2010) | 5B | None |
| 21 | *Halopiger djelfimassiliensis* | IIH2 | 1293047 | CBMA0000 00000 | None | 3560 | Hassani *et al.* (2013) | 5C | gen. nov. |
| 22 | *Halopiger goleimassiliensis* | IIH3 | 1293048 | CBMB0000 00000 | None | 3705 | Ikram *et al.* (2014) | 5C | gen. nov. |
| 23 | '*Natrarchaeobius chitinivorans*' | AArch4<sup>T</sup> | 1679083 | REGA00000 000 | UP000282323 | 4237 | Sorokin *et al.* (2019a) | 5D | None |
| 24 | '*Natrarchaeobius chitinivorans*' | AArch7 | 1679083 | REFZ00000 000 | UP000281431 | 4321 | None | 5D | gen. nov. |
| 25 | '*Natrarchaeobius halalkaliphilus*' | AArch-SI<sup>T</sup> | 1679091 | REFY00000 000 | UP000273828 | 3330 | Sorokin *et al.* (2019a) | 5D | gen. nov. |
| 26 | *Natronolimnobius aegyptiacus* | JW/NM-HA 15<sup>T</sup> | 745377 | CP019893 | UP000250088 | 3626 | Zhao *et al.* (2018) | 5D | gen. nov. |
| 27 | *Natronolimnobius sulfurireducens* | AArc1<sup>T</sup> | 2044521 | CP024047 | UP000258707 | 3466 | Sorokin *et al.* (2019b) | 5D | gen. nov. along with serial number 28 |
| 28 | *Natronolimnobius sulfurireducens* | AArc-Mg | 2086290 | CP027033 | UP000258613 | 3528 | Sorokin *et al.* (2019b) | 5D |  |
| 29 | *Halobiforma haloterrestris* | DSM 13078<sup>T</sup> | 148448 | FOKW0000 0000 | UP000199161 | 4334 | Hezayen *et al.* (2002) | 5E | None |
| 30 | *Halobiforma lacisalsi* | AJ5<sup>T</sup> | 358396 | CP019285 | UP000186547 | 3913 | Xu *et al.* (2005b) | 5E | None |
| 31 | *Halobiforma lacisalsi* | AJ5<sup>T</sup> | 358396 | AOLZ01000 000 | UP000011555 | 4151 | Xu *et al.* (2005b) | 5E | None |
| 32 | *Halobiforma nitratreducens* | JCM 10879<sup>T</sup> | 1227454 | AOMA0000 0000 | UP000011607 | 3534 | Hezayen *et al.* (2002) | 5E | gen. nov. |
| 33 | *Natronobacterium gregoryi* | ATCC 43098<sup>T</sup> | 797304 | AOIC01000 000 | UP000011613 | 3655 | Tindall *et al.* (1984) | 5F | None |
| 34 | *Natronobacterium gregoryi* | ATCC 43098<sup>T</sup> | 797304 | PKKI01000 000 | UP000234484 | 3568 | Tindall *et al.* (1984) | 5F | None |
| 35 | *Natronobacterium gregoryi* | SP2<sup>T</sup> | 44930 | FORO01000 000 | UP000182829 | 3713 | Tindall *et al.* (1984) | 5F | None |
| 36 | *Natronobacterium gregoryi* | ATCC 43098<sup>T</sup> | 797304 | CP003377 | UP000010468 | 3624 | Tindall *et al.* (1984) | 5F | None |
| 37 | *Natronobacterium texcoconense* | DSM 24767<sup>T</sup> | 1095778 | FNLC01000 000 | UP000198848 | 4046 | Ruiz-Romero *et al.* (2013a) | 5F | gen. nov. |
| 38 | *Natronorubrum bangense* | JCM 10635<sup>T</sup> | 1227500 | AOHY0100 0000 | UP000011690 | 3979 | Xu *et al.* (1999) | 6A | None |
| 39 | *Natronorubrum sulfidifaciens* | JCM 14089<sup>T</sup> | 1230460 | AOHX0100 0000 | UP000011661 | 3428 | Cui *et al.* (2007) | 6A | None |
| 40 | '*Natronorubrum thiooxidans*' | HArc- | 308853 | FTNR01000 000 | UP000185936 | 4231 | None | 6A | None |
| 41 | *Natronorubrum tibetense* | GA33<sup>T</sup> | 1114856 | AOHW0100 0000 | UP000011599 | 4671 | Xu *et al.* (1999) | 6A | gen. nov. along with serial number 42 |
| 42 | *Natronorubrum texcoconense* | B4<sup>T</sup> | 1095776 | FNFE01000 000 | UP000198882 | 4464 | Ruiz-Romero *et al.* (2013b) | 6A |  |
| 43 | *Natronorubrum sediminis* | CGMCC 1.8981<sup>T</sup> | 640943 | FNWL0100 0000 | UP000199112 | 3617 | Gutiérrez *et al.* (2010) | 6A | gen. nov. along with serial number 44/45 |
| 44 | *Haloterrigena daqingensis* | JX313<sup>T</sup> | 588898 | CP019327 | UP000187321 | 3195 | Wang *et al.* (2010) | 6A |  |
| 45 | *Haloterrigena daqingensis* | CGMCC 1.8909<sup>T</sup> | 588898 | FTNP01000 000 | UP000185687 | 3795 | Wang *et al.* (2010) | 6A |  |
| 46 | *Natronolimnobius innermongolicus* | JCM 12255<sup>T</sup> | 1227499 | AOHZ0100 0000 | UP000011602 | 4400 | Itoh *et al.* (2005) | 6A | gen. nov. |
| 47 | *Haloterrigena turkmenica* | ATCC 51198<sup>T</sup> | 543526 | CP001860 | UP0000001903 | 3739 | Ventosa *et al.* (1999) | 6B | None |
| 48 | *Haloterrigena salina* | JCM 13891<sup>T</sup> | 1227488 | AOIS01000 000 | UP000011657 | 4563 | Gutiérrez *et al.* (2008) | 6B | None |
| 49 | *Halopiger salifodinae* | CGMCC 1.12284<sup>T</sup> | 1202768 | FOIS010000 00 | UP000183275 | 4090 | Zhang *et al.* (2013) | 6C | gen. nov. |
| 50 | *Haloterrigena limicola* | JCM 13563<sup>T</sup> | 1230457 | AOIT01000 000 | UP000011615 | 3519 | Cui *et al.* (2006) | 6D | gen. nov. along with serial numbers 51, 52, and 53 |
| 51 | *Haloterrigena hispanica* | DSM 18328<sup>T</sup> | 392421 | SHMP01000 000 | UP000291097 | 4092 | Romano *et al.* (2007) | 6D |  |
| 52 | *Haloterrigena* sp. | CDM_6 | 392421 | FOIC01000 000 | UP000199320 | 4122 | None | 6D |  |
| 53 | *Haloterrigena* sp. | H1 | 2552943 | SMZK0000 0000 | None | 3974 | None | 6D |  |
| 54 | *Natrinema pellirubrum* | DSM 15624<sup>T</sup> | 797303 | AOIE01000 000 | UP000011593 | 4176 | McGenity *et al.* (1998) | 6E | None |
| 55 | *Natrinema pellirubrum* | DSM 15624<sup>T</sup> | 797303 | CP003372 | UP000010843 | 3653 | McGenity *et al.* (1998) | 6E | None |
| 56 | *Haloterrigena thermotolerans* | DSM 11522<sup>T</sup> | 1227489 | AOIR01000000 | UP000011594 | 3858 | Montalvo-Rodríguez *et al.* (2000) | 6E | Merge with *Nn. pellirubrum* |
| 57 | *Haloterrigena jeotgali* | A29<sup>T</sup> | 413811 | CP031303 | None | 3,550 | Roh *et al.* (2009) | 6E | Merge with *Nn. pellirubrum* |
| 58 | *Haloterrigena saccharevitans* | AB14<sup>T</sup> | 301967 | LWLN01000000 | UP000189370 | 3699 | Xu *et al.* (2005c) | 6E | comb. nov. |
| 59 | *Haloterrigena mahii* | H13<sup>T</sup> | 1416969 | JHUT02000000 | UP000053318 | 3790 | Ding *et al.* (2017) | 6E | comb. nov. |
| 60 | *Natrinema versiforme* | JCM 10478<sup>T</sup> | 1227496 | AOID01000000 | UP000011632 | 4160 | Xin *et al.* (2000) | 6E | gen. nov. |
| 61 | '*Natrinema thermophila*' | CBA1119 | 1608465 | PDBS01000000 | UP000223005 | 4602 | Kim *et al.* (2018) | 6E | gen. nov. |
| 62 | *Natrinema salaciae* | DSM 25055<sup>T</sup> | 1186196 | FOFD01000000 | UP000199114 | 4688 | Albuquerque *et al.* (2012) | 6E | gen. nov. |
| 63 | *Natrinema ejinorensense* | JCM 13890<sup>T</sup> | 373386 | NXNI01000000 | UP000219689 | 4191 | Castillo *et al.* (2006c) | 6E | gen. nov. |
| 64 | *Natrinema gari* | JCM 14663<sup>T</sup> | 1230459 | AOIJ01000000 | UP000011592 | 4056 | Tapingkae *et al.* (2008) | 6E | gen. nov. along with serial numbers 65, 66, and 67 |
| 65 | *Natrinema* sp. | J7-2 | 406552 | CP003412 | UP0000006507 | 4233 | Feng *et al.* (2012) | 6E |  |
| 66 | *Natrinema pallidum* | DSM 3751<sup>T</sup> | 1227495 | AOII01000000 | UP000011618 | 3844 | McGenity *et al.* (1998) | 6E |  |
| 67 | *Natrinema altunense* | JCM 12890<sup>T</sup> | 1227494 | AOIK01000000 | UP000011511 | 3732 | Xu *et al.* (2005a) | 6E |  |

### Phylogenetic analysis using CVTree3

Alignment-free sequence comparison methods offer several advantages over alignment-based methods, and are particularly useful in the analysis of large datasets (Zielezinski *et al*., 2017). CVTree3 is an alignment- and parameter-free method that relies on the oligopeptide content (K-tuple length) of conserved proteins to deduce evolutionary relatedness (Zuo and Hao 2015), and has been used previously for the phylogenetic characterization of archaeal taxa at and above the order level (Zuo *et al*., 2015). Phylogenetic analysis using CVTree3 was performed as described previously (Viswanathan *et al*., 2017; Siddaramappa *et al*., 2019). Briefly, the deduced proteome sequences downloaded from UniProt were saved as multifasta files with the extension .faa. The multifasta files from different species/strains were uploaded on to the CVTree3 web server (http://tlife.fudan.edu.cn/cvtree/cvtree/) and analyzed by selecting all available K-tuple length options (from 3 to 9). Because the best K-values for prokaryotes were shown to be 5–6 (Zuo *et al*., 2015; Zuo and Hao 2015), the deduced proteome sequence-based tree was visualized at K = 6. The output from CVTree3 was saved as a Newick file, and the tree was rendered using the Interactive Tree Of Life (https://itol.embl.de/) web server version 4 (Letunic and Bork 2019).

## RESULTS AND DISCUSSION

### Phylogenetic insights based on deduced proteome sequences

The number of predicted proteins among the 67 genome sequences varied from 2711– 4688 (Table 1). Six major clades were conspicuous in the deduced proteome sequence-based phylogenetic tree, and majority of the species/strains were located on clades 5 and 6 (Fig. 1). To further understand the phylogenetic relationships among these species/strains, the criterion proposed by Chun *et al*. (2018) was used. If the ANI was ≥95%, then the strain was inferred not to represent a novel species. Although Qin *et al*. (2014) opined that ANI comparisons may not be suitable for genus delimitation, phylogenetic analysis combined with ANI comparisons could be useful to identify novel genera. In this study, if the ANI was ≥85%, then the strain was inferred not to represent a novel genus. For example, *Halovivax asiaticus* and *Halovivax ruber* were co-located on a separate branch in the phylogenetic tree (Fig. 1). The ANI value between *Hv. asiaticus*, the type species of the genus, and *Hv. ruber* was 92.41%. The AAI value between these archaea was 92.99%. Therefore, the proposals that *Halovivax* is a novel genus (Castillo *et al*., 2006a), and that *Hv. ruber* is a novel species of this genus (Castillo *et al*., 2007), are valid. Likewise, *Halostagnicola larsenii*, *Halostagnicola kamekurae*, and *Halostagnicola* sp. A56 were co-located on a separate branch in the phylogenetic tree (Fig. 1). The ANI/AAI values between *Hs. larsenii*, the type species of the genus, and *Hs. kamekurae* and *Halostagnicola* sp. A56 were 87.16%/86.61% and 86.88%/86.84%, respectively. Therefore, the proposals that *Halostagnicola* is a novel genus (Castillo *et al*., 2006b), and that *Hs. kamekurae* is a novel species of this genus (Nagaoka *et al*., 2010), are valid. It is also likely that *Halostagnicola* sp. A56, whose ANI/AAI value with *Hs. kamekurae* was 92.10%/91.64%, represents a novel species that is yet to be characterized and named. Furthermore, *Natronococcus occultus*, *Natronococcus amylolyticus*, and *Natronococcus jeotgali* were co-located on a separate branch in the phylogenetic tree (Fig. 1). The ANI/AAI values between *Nc. occultus*, the type species of the genus, and *Nc. amylolyticus* and *Nc. jeotgali* were 85.82%/84.69% and 87.01%/84.22%, respectively. Hence, the proposals that *Natronococcus* is a novel genus (Tindall *et al*., 1984), and that *Nc. amylolyticus* and *Nc. jeotgali* are novel species of this genus (Kanal *et al*., 1995; Roh *et al*., 2007), are valid.

**Fig. 1.**
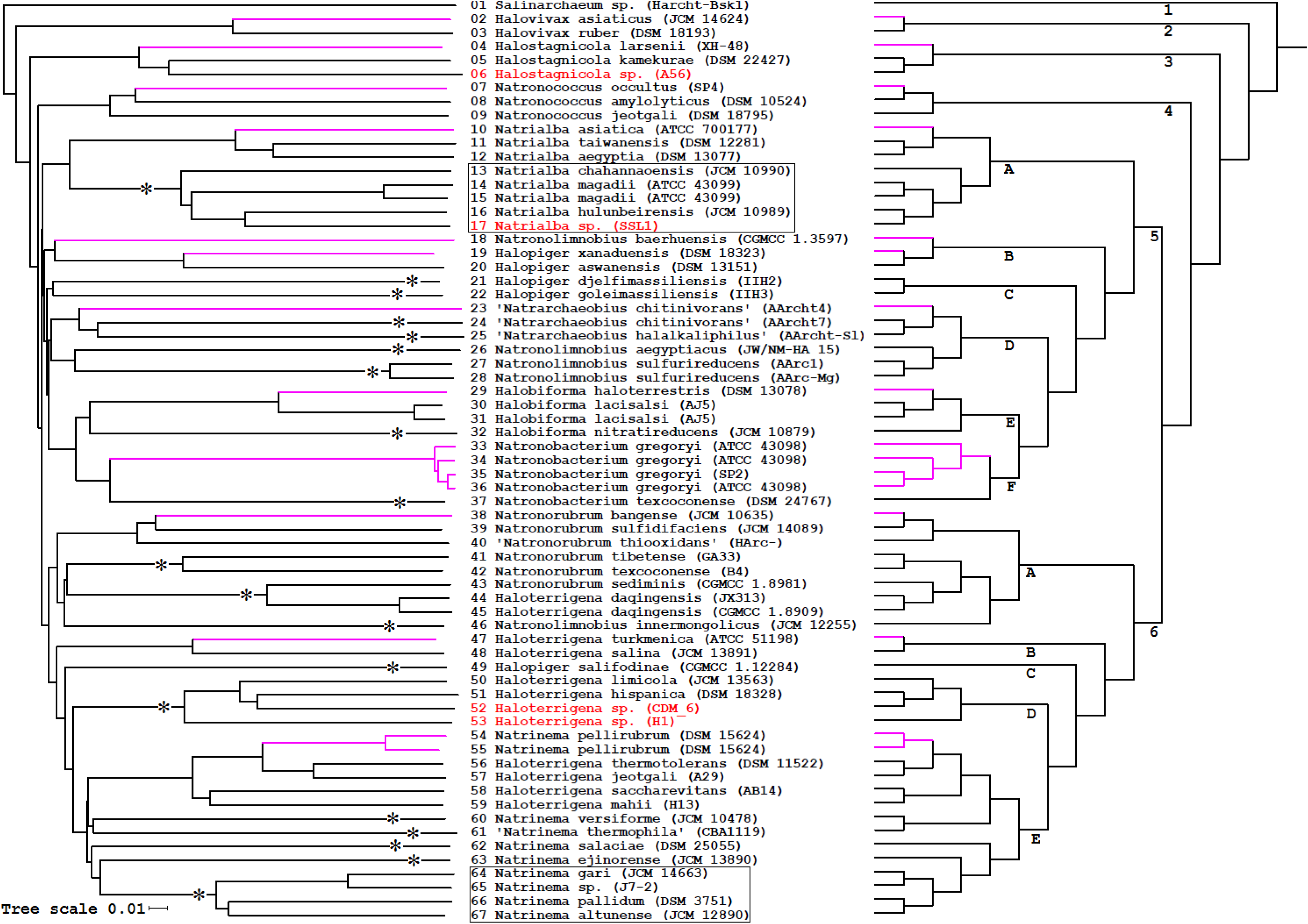
Phylogenetic tree based on deduced proteome sequences. The tree was constructed using the neighbor-joining method by the web server CVTree3, and visualized at K = 6. The proteome of *Halalkalicoccus jeotgali* B3 was used as the outgroup, which does not appear in the figure. The scale bar below the tree on the left allows the estimation of branch lengths, and numbers at the end of each branch refer to the serial numbers in Table 1. Strains likely to represent novel species are shown using red text, and asterisks indicate strains likely to represent novel genera. *Natrialba* and *Natrinema* species likely to represent novel genera are boxed. The tree on the right was drawn without taking branch lengths into consideration, and numbers 1–6 indicate the major clades (subclades A–F of clade 5, and subclades A–E of clade 6, are also marked). Pink lines indicate type species with validly/effectively published names (n = 12).

### The genus *Natrialba* and a potentially novel genus within subclade 5A

The novel genus *Natrialba* containing a single species was proposed by Kamekura and Dyall-Smith (1995). Since then, the names of five species of *Natrialba* have been validly published. As of July 2019, the genome sequences of all six species of *Natrialba* have been sequenced, with *Natrialba magadii* ATCC 43099^T^ being sequenced twice (Table 1). Subclade 5A in the phylogenetic tree (Fig. 1) contained two distinct branches, each having three species of *Natrialba*. The ANI/AAI value between *Natrialba taiwanensis* and *Natrialba aegyptia* was 94.83%/95.00%. Since the DNA–DNA hybridization level between *Na. taiwanensis* and *Na*. *aegyptia* was reported to be 66% (Hezayen *et al*., 2001), it is likely that these two species are very closely related. The DNA–DNA hybridization level between *Natrialba asiatica*, the type species of the genus, and *Na. taiwanensis* was reported to be 55.8% (Hezayen *et al*., 2001), indicating that they are two different species. This is supported by the fact that the ANI/AAI value between these archaea was 93.12%/92.95%.

Furthermore, the ANI/AAI value between *Natrialba chahannaoensis* and *Na. magadii* was 88.84%/88.17%, the ANI/AAI value between *Na. magadii* and *Natrialba hulunbeirensis* was 89.40%/89.61%, and the ANI/AAI value between *Na. chahannaoensis* and *Na*. *hulunbeirensis* was 88.78%/89.90%. This result confirms the proposal by Xu *et al*. (2001) that *Na. chahannaoensis* and *Na. hulunbeirensis* are distinct from each other, and from *Na. magadii*. However, the ANI/AAI values between *Na. asiatica* and *Na. chahannaoensis*, *Na. magadii*, *Na. hulunbeirensis*, and *Natrialba* sp. SSL1 were 82.17%/76.00%, 82.58%/76.21%, 82.55%/76.30%, and 82.75%/75.75%. Interestingly, Hezayen *et al*. (2001) had reported that the 16S rDNA sequence similarity between *Na. asiatica* and *Na. magadii* was <96%, and Xu *et al*. (2001) had reported that the 16S rDNA sequence similarity between *Na. chahannaoensis* (or *Na. hulunbeirensis*) and *Na. asiatica* was <96%. Based on these results, it is likely that *Na. chahannaoensis*, *Na. magadii*, and *Na. hulunbeirensis* together represent a novel genus that is different from the one represented together by *Na. asiatica*, *Na. taiwanensis*, and *Na. aegyptia*. Since the ANI/AAI value between *Na. hulunbeirensis* and *Natrialba* sp. SSL1 was 91.41%/92.71%, the latter represents a novel species that is yet to be characterized and named.

### The genera *Natronolimnobius* and *Halopiger* within subclade 5B

Subclade 5B in the phylogenetic tree (Fig. 1) contained *Natronolimnobius baerhuensis* on a deep branch, and *Halopiger xanaduensis* and *Halopiger aswanensis* on a separate branch. The ANI/AAI values between *Nl. baerhuensis*, the type species of the genus, and *Hp. xanaduensis* and *Hp. aswanensis* were 81.09%/74.38% and 80.89%/74.03%, respectively. Therefore, the genera *Natronolimnobius* and *Halopiger*, as proposed by Itoh *et al*. (2005) and Gutiérrez *et al*. (2007), respectively, are sufficiently different from each other. Since the ANI/AAI value between *Hp. xanaduensis* and *Hp. aswanensis* was 89.60%/90.93%, the proposal by Hezayen *et al*. (2010) that the latter represents a novel species is valid.

### Two potentially novel genera within subclade 5C

Subclade 5C in the phylogenetic tree (Fig. 1) contained *Halopiger djelfimassiliensis* and *Halopiger goleimassiliensis*, each located on a deep branch. These two archaea were isolated from the Djelfa and Ghardaïa regions, respectively, of Algeria (Hassani *et al*., 2013; Ikram *et al*., 2014). It was reported that the 16S rDNA sequence similarity between *Hp. djelfimassiliensis* (or *Hp. goleimassiliensis*) and *Hp. xanaduensis* was ~96% (Hassani *et al*., 2013; Ikram *et al*., 2014). The ANI/AAI values between *Hp. xanaduensis*, the type species of the genus, and *Hp. djelfimassiliensis* and *Hp. goleimassiliensis* were 82.43%/75.17% and 83.03%/73.52%, respectively, indicating that the two archaea within subclade 5C are not members of the genus *Halopiger*. Furthermore, the ANI/AAI value between *Hp. djelfimassiliensis* and *Hp. goleimassiliensis* was 82.00%/74.05%. Therefore, it appears that subclade 5C represents two distinct genera of the family *Natrialbaceae*.

### Five potentially novel genera within subclade 5D

One of the branches within subclade 5D contained ‘*Natrarchaeobius chitinivorans*’ and ‘*Natrarchaeobius halalkaliphilus*’ (Sorokin *et al*., 2019a), which are yet to be validly published. The ANI/AAI value between ‘*Nb. chitinivorans*’ Aarcht4^T^ and ‘*Nb. chitinivorans*’ Aarcht7 was 83.56%/76.74%, the ANI/AAI value between ‘*Nb. chitinivorans*’ Aarcht7 and ‘*Nb. halalkaliphilus*’ was 82.40%/78.83%, and the ANI/AAI value between ‘*Nb. chitinivorans*’ Aarcht4^T^ and ‘*Nb. halalkaliphilus*’ was 82.03%/75.38%. Therefore, it is likely each of these three archaea represent novel genera of the family *Natrialbaceae*.

The other branch within subclade 5D contained *Natronolimnobius aegyptiacus* and *Natronolimnobius sulfurireducens*. The ANI/AAI values between *Nl. baerhuensis*, the type species of the genus, and *Nl. aegyptiacus* and *Nl. sulfurireducens* were 80.09%/69.98% and 79.94%/71.17%, respectively. Furthermore, the ANI/AAI value between *Nl. aegyptiacus* and *Nl. sulfurireducens* was 81.87%/75.46%. Therefore, not only are *Nl. aegyptiacus* (Zhao *et al*., 2018) and *Nl. sulfurireducens* (Sorokin *et al*., 2019b) misidentified, but also appear to represent two novel genera within the family *Natrialbaceae*.

### Two potentially novel genera within subclades 5E and 5F

The inclusion of *Halobiforma nitratireducens* within the genus *Halobiforma* by Hezayen *et al*. (2002) was not unambiguous due to the polyphyletic nature of the strains used in the analyses. Since the ANI/AAI value between *Halobiforma haloterrestris* and *Hb. nitratireducens* was 83.63%/79.15%, it is likely that the latter represents a novel genus by itself within subclade 5E. Furthermore, Xu *et al*. (2005b) reported that the 16S rDNA gene sequences of *Halobiforma lacisalsi* and *Hb. haloterrestris* had 99.0% similarity. The ANI/AAI value between *Hb. haloterrestris* and *Hb. lacisalsi* was 94.62%/95.07%, confirming that these two species are very closely related.

Within subclade 5F, the ANI/AAI value between *Natronobacterium gregoryi* and *Natronobacterium texcoconense* was 84.75%/79.15%. Although the 16S rDNA gene sequences of these archaea were reported to have 97.3% similarity, the DNA–DNA hybridization level between them was only 32.3% (Ruiz-Romero *et al*., 2013a). Notably, *Nb. texcoconense* was oxidase negative, and the optimum NaCl concentration at which it could grow was different from that of *Nb. gregoryi* (Ruiz-Romero *et al*., 2013a). The optimum pH at which these archaea could grow was also different (Ruiz-Romero *et al*., 2013a). Based on these results, it is proposed that *Nb. texcoconense* represents yet another novel genus.

### Three potentially novel genera within subclade 6A

The novel genus *Natronorubrum* was proposed by Xu *et al*. (1999) to include *Natronorubrum bangense* and *Natronorubrurn tibetense*, with the former as the type species. Since then, the names of four species of *Natranorubrum* have been validly published. As of July 2019, the genome sequences of five species of *Natronorubrum* have been sequenced (Table 1). Within subclade 6A, *Nr. bangense* and *Natronorubrum sulfidifaciens* were co-located, and the ANI/AAI value between these two species was 88.49%/87.12%. Therefore the proposal by Cui *et al*. (2007) that *Nr. sulfidifaciens* is a novel species of the genus is valid. Although the name *’Natronorubrum thiooxidans’* has not yet been effectively/validly published, this species was co-located with *Nr. bangense* and *Nr. sulfidifaciens* in the phylogenetic tree (Fig. 1). Furthermore, ANI/AAI values between *’Nr. thiooxidans’* and the other two archaea in the cluster were 86.67%/85.48% and 86.66%/86.06%. Therefore, *Nr. bangense*, *Nr. sulfidifaciens*, and *’Nr. thiooxidans’* are distinct from each other.

Ruiz-Romero *et al*. (2013b) proposed *Natronorubrum texcoconense* as a novel species based on chemotaxonomic studies and the comparison of 16S rDNA gene sequences. It was reported that the 16S rDNA gene sequence of *Nr. texcoconense* had 96.28%, 95.06%, and 94.98% similarity to the 16S rDNA gene sequences of *Nr. tibetense*, *Nr. sulfidifaciens*, and *Natronorubrum sediminis*, respectively. Not surprisingly, in the proteome sequence-based tree, *Nr. texcoconense* clustered with *Nr. tibetense*, and away from *Nr. sulfidifaciens* (Fig. 1). Furthermore, the ANI/AAI value between *Nr. texcoconense* and *Nr. tibetense* was 89.86%/90.63%. Since the ANI/AAI value between *Nr. bangense* and *Nr. texcoconense* was 81.71%/76.58%, and the ANI/AAI value between *Nr. bangense* and *Nr. tibetense* was 82.27%/ 77.00%, it is likely that *Nr. texcoconense* and *Nr. tibetense* represent a novel genus within subclade 6A.

The co-location of *Nr. texcoconense* and *Nr. tibetense* with *Nr. sedeminis* in the phylogenetic tree (Fig. 1) insinuated that they are related. However, the ANI/AAI values between *Nr. sedeminis* and *Nr. texcoconense*, *Nr. tibetense*, and *Nr. bangense* were 81.62%/77.20%, 81.50%/76.78%, and 80.49%/74.28%, respectively. Therefore, it is likely that *Nr. sedeminis* also represents a novel genus within subclade 6E. The co-location of *Haloterrigena daqingensis* (Wang *et al*., 2010) with *Nr. sedeminis* (Fig. 1), and not with *Haloterrigena turkmenica*, raised questions about its taxonomic status. BLASTN analysis showed that the 16S rDNA gene sequence of *Ht. daqingensis* had 99.80%, 95.87%, 95.46%, and 96.68% identity to the 16S rDNA gene sequences of *Nr. sediminis*, *Nr. sulfidifaciens*, *Nr. bangense*, and *Ht. turkmenica*. Likewise, the 16S rDNA gene sequence of *Nr. sediminis* had 96.07%, 95.67%, and 96.75% identity to the 16S rDNA gene sequences of *Nr. sulfidifaciens*, *Nr. bangense*, and *Ht. turkmenica*. Based on these results, it appears that *Ht. daqingensis* and *Nr. sediminis* are more closely related to *Ht. turkmenica*. However, the ANI/AAI value between *Ht. turkmenica* and *Ht. daqingensis* was 81.54%/74.90%, the ANI/AAI value between *Ht. turkmenica* and *Nr. sedeminis* was 81.25%/74.13%, and the ANI/AAI value between *Nr. sedeminis* and *Ht. daqingensis* was 93.35%/95.04%. Therefore, it is proposed that *Ht. daqingensis* be included in a novel genus along with *Nr. sedeminis*.

The location of *Natronolimnobius innermongolicus* within subclade 6A (Fig. 1), and not with *Natronolimnobius baerhuensis*, also raised questions about its taxonomic status. The ANI/AAI value between *Nl. baerhuensis* and *Nl. innermongolicus* was 80.83%/71.35%, and BLASTN analysis showed that the 16S rDNA gene sequences of these archaea had only 96.04% identity. Furthermore, the ANI/AAI values between *Nl. innermongolicus* and *Nr. bangense*, *Nr. texcoconense*, *Nr. tibetense*, and *Nr. sedeminis* were 81.70% /74.62%, 82.42%/76.10%, 82.26%/75.92%, and 81.66%/75.55%, respectively. Therefore, it is likely that *Nl. innermongolicus* represents yet another novel genus within this subclade.

### Two potentially novel genera within subclades 6C and 6D

*Halopiger salifodinae* was proposed as a novel species of the genus *Halopiger* by Zhang *et al*. (2013), who reported that the 16S rDNA gene sequences of this archaeon and *Hp. xanaduensis* had 95.8% similarity. However, *Hp. salifodinae* was located away from *Hp. xanaduensis*, which is the type species of the genus, on a separate deep branch (Fig. 1). While the ANI/AAI value between *Hp. xanaduensis* and *Hp. aswanensis* was 89.60%/90.93%, the ANI/AAI value between *Hp. xanaduensis* and *Hp. salifodinae* was 83.22%/74.94%. Therefore, it appears that *Hp. salifodinae* (subclade 6C) is only distantly related to members of the genus *Halopiger*, and represents an yet to be characterized novel genus.

*Haloterrigena limicola*, *Haloterrigena hispanica*, *Haloterrigena* sp. CDM_6, and *Haloterrigena* sp. H1 clustered away from *Ht. turkmenica* on a separate branch (Fig. 1). While the 16S rDNA gene sequences of *Ht. limicola* and *Ht. turkmenica* had 93.9% similarity (Cui *et al*., 2006), the 16S rDNA gene sequences of *Ht. hispanica* and *Ht. turkmenica* had 95.5% similarity (Romano *et al*., 2007), indicating that *Ht. limicola* and *Ht. hispanica* are distantly related to *Ht. turkmenica*. However, the 16S rDNA gene sequences of *Ht. hispanica* and *Ht. limicola* had 98.9% similarity (Romano *et al*., 2007), suggesting that they are closely related to each other. Although the ANI/AAI value between *Ht. turkmenica* and *Haloterrigena salina* (co-located in subclade 6B of Fig. 1) was 91.42%/91.28%, the ANI/AAI values between *Ht*. *turkmenica* and the four strains in subclade 6D were 82.32%/74.77%, 82.46%/74.13%, 82.15%/74.37%, and 82.46%/74.66%, respectively. Based on these results, it is proposed that subclade 6D represents a novel genus of *Natrialbaceae*. Furthermore, the ANI/AAI value between *Ht. limicola* and *Ht. hispanica* was 93.82%/94.35%, and the ANI/AAI value between *Ht. limicola* and *Haloterrigena* sp. CDM_6 was 92.46%/93.64%. Since the ANI/AAI value between *Ht. hispanica* and *Haloterrigena* sp. CDM_6 was 93.64%/94.50%, the latter represents a species that is sufficiently novel. Likewise, *Haloterrigena* sp. H1 represents a novel species within subclade 6D because the ANI/AAI value between *Ht. limicola* and this strain was 88.83%/90.01%.

### Genus *Natrinema* and five potentially novel genera within subclade 6E

The novel genus *Natrinema* was proposed by McGenity *et al*. (1998) to include *Natrinema pellirubrum* and *Natrinerna pallidum*, with the former as the type species. Since then, the names of six species of *Natrinema* have been validly published (Table 1). As of July 2019, the genome sequences of all eight species of *Natrinema* have been sequenced, with *Natrinema pellirubrum* DSM 15624^T^ being sequenced twice (Table 1). Subclade 6E in Fig. 1 contained all the eight *Natrinema* spp. and four *Haloterrigena* spp. Roh *et al*. (2009) reported that the 16S rDNA gene sequences of *Haloterrigena thermotolerans* and *Haloterrigena jeotgali* had 99% similarity, indicating that they are very closely related. Since the ANI/AAI value between these archaea was 97.75%/96.97%, it is likely that they are not two species. Furthermore, despite sufficient differences in phenotypic properties, Montalvo-Rodríguez *et al*. (2000) assigned *Ht. thermotolerans* to the genus *Haloterrigena*. Although 16S rDNA gene sequence analysis indicated that *Ht. thermotolerans* was related to members of the genera *Haloterrigena* and *Natrinema*, the DNA–DNA hybridization level between *Ht. thermotolerans* and *Ht. turkmenica* was reported to be only 48% (Montalvo-Rodríguez *et al*., 2000). Xu *et al*. (2005c) reported that the 16S rDNA gene sequences of *Haloterrigena saccharevitans* and *Ht. thermotolerans* had 98.6% similarity, and that the 16S rDNA gene sequences of *Ht. saccharevitans* and *Ht. turkmenica* had only 96% similarity. Likewise, Ding *et al*. (2017) reported that the 16S rDNA gene sequences of *Haloterrigena mahii* and *Ht. thermotolerans* had 99.11% similarity. However, the 16S rDNA gene sequences of *Ht. mahii* and *Ht. turkmenica* had only 97% identity, while the 16S rDNA gene sequences of *Ht. mahii* and *Nn. pellirubrum* had 99.32% identity. Whole genome analyses showed that the ANI/AAI values between *Ht. turkmenica*, the type species of the genus, and *Ht. thermotolerans*, *Ht. jeotgali*, *Ht. saccharevitans*, and *Ht. mahii* were 82.90%/75.59%, 83.12%/75.50%, 83.26%/75.39%, 83.26%/75.42%, respectively. These results clearly indicate that the four *Haloterrigena* spp. in Subclade 6E are only distantly related to *Ht. turkmenica*. Since the ANI/AAI value between *Nn. pellirubrum* and *Ht. thermotolerans* was 95.60%/95.47% (and the ANI/AAI value between *Nn. pellirubrum* and *Ht. jeotgali* was 95.31%/95.14%), it is proposed to merge *Ht. thermotolerans* and *Ht. jeotgali* with *Nn. pellirubrum*. The ANI/AAI value between *Ht. saccharevitans* and *Ht. mahii* was 92.62%/92.12%, indicating that they are two different species. Because the ANI/AAI values between these two species and *Nn. pellirubrum* were 90.51%/90.24% and 91.67%/92.09%, respectively, it is proposed that they be transferred to the genus *Natrinema* as *Natrinema saccharevitans* comb. nov. and *Natrinema mahii* comb. nov.

Within subclade 6E, *Natrinema versiforme* and ‘*Natrinema thermophila*’ were located on deep branches. The 16S rDNA gene sequences of these archaea had 96.09% identity, and the ANI/AAI value between them was 84.48%/80.87%. Furthermore, the 16S rDNA gene sequence of *Nn. pellirubrum* had 97.22% and 96.36% identity, respectively, to those of *Nn. versiforme* and ‘*Nn. thermophila*’. The ANI/AAI values between *Nn. pellirubrum* and these two archaea were 84.12%/80.96% and 83.44%/79.86%. Therefore, it is likely that *Nn. versiforme* and ‘*Nn. thermophila*’ each represent a novel genus.

*Natrinema salaciae* and *Natrinema ejinorense* were also located on deep branches away from *Nn. pellirubrum*. The 16S rDNA gene sequences of these archaea had 97.60% identity, and the ANI/AAI value between them was 84.78%/81.44%. Furthermore, the 16S rDNA gene sequence of *Nn. pellirubrum* had 96.65% and 96.13% identity, respectively, to those of *Nn. salaciae* and *Nn. ejinorense*. The ANI/AAI values between *Nn. pellirubrum* and these two archaea were 83.47%/80.29% and 83.78%/80.26%, suggesting that *Nn. salaciae* and *Nn. ejinorense* each represent a novel genus.

Feng *et al*. (2012) reported that *Natrinema* sp. J7-2 and *Natrinema gari* are closely related. Not surprisingly, the ANI/AAI value between them was 98.79%/ 98.11%, further confirming that the former is yet another strain of *Nn. gari*. Since the ANI/AAI value between *Natrinema pallidum* and *Natrinema altunense* was 93.26%/93.17%, the proposal by Xu *et al*. (2005a) that the latter is a novel species is valid. The ANI/AAI values between *Nn. gari* and these two archaea were 92.16%/91.31% and 92.80%/92.66%, suggesting that they are closely related, but represent distinct species. However, the ANI/AAI values between *Nn. ejinorense* and *Nn. gari*, *Nn. pallidum*, and *Nn. altunense* were 84.76%/80.89%, 84.72%/81.52%, and 85.21%/82.38%, respectively. Therefore, the entire cluster containing *Nn. gari*, *Nn. pallidum*, and *Nn. altunense* may represent a novel genus.

## ACKNOWLEDGEMENTS

This work was supported by facilities and funds provided by the Department of Information Technology, Biotechnology, and Science & Technology of the Government of Karnataka to the Institute of Bioinformatics and Applied Biotechnology (IBAB). Research infrastructure at IBAB is made possible by the Department of Science & Technology of the Government of India through a program called “Fund for Improvement of S&T infrastructure in universities & higher educational institutions” (*DST-FIST*: Level 0 from 2006 to 2011, and Level 1 from 2012 to 2017). The author is grateful to the global scientific community for freely sharing the genome sequences of haloalkaliphilic archaea in the public databases.

